# Sensitive and specific post-call filtering of genetic variants in xenograft and primary tumors

**DOI:** 10.1101/187468

**Authors:** Brian K Mannakee, Uthra Balaji, Agnieszka K. Witkiewicz, Ryan N. Gutenkunst, Erik S. Knudsen

## Abstract

**Motivation:** Tumor genome sequencing offers great promise for guiding research and therapy, but spurious variant calls can arise from multiple sources. Mouse contamination can generate many spurious calls when sequencing patient-derived xenografts (PDXs). Paralogous genome sequences can also generate spurious calls when sequencing any tumor. We developed a BLAST-based algorithm, MAPEX, to identify and filter out spurious calls from both these sources.

**Results:** When calling variants from xenografts, MAPEX has similar sensitivity and specificity to more complex algorithms. When applied to any tumor, MAPEX also automatically flags calls that potentially arise from paralogous sequences. Our implementation, mapexr, runs quickly and easily on a desktocomputer. MAPEX is thus a useful addition to almost any pipeline for calling genetic variants in tumors.

## 1 Introduction

Molecular characterization of tumors is an important tool in cancer research, and the large-scale sequencing of cancer genomes has led to a deeper understanding of many aspects of the biology of cancer [Stratton MR, 2011]. It is now common to sequence tumors from large cohorts of patients, as well as patient-derived xenograft (PDX) models from individual patients. Such sequencing enables identification of mutational signatures [Alexandrov et al., 2013], functionally important variants [Ding et al., 2012] and evolutionary history of the tumor [Carter et al., 2012, Nik-Zainal et al., 2012]. These genetic features are relevant in evaluating etiological mechanisms [Yachida et al., 2010], prognostic subtypes [Park et al., 2010, Shah et al., 2009], and acquired therapeutic resistance [Witkiewicz et al., 2015]. All these applications of tumor sequencing depend on sensitive and specific characterization of low-frequency mutations, and as a result may be biased by spurious variant calls. Here we focus on two specific sources of spurious calls, mouse cell contamination in PDX tumors and mis-alignment of paralogous sequences.

PDX models serve as avatars for individual patient tumors when studying intra-tumor heterogeneity and metastasis and when screening anti-cancer compounds [Allaway et al., 2016, Bruna et al., 2016,Dawson et al., 2012, Day et al., 2015, Knudsen et al., 2017]. The primary difficulty in sequencing these models is that mouse stroma is present in all PDX tumors. The high genetic similarity between mouse and human then causes bias when variants are called using bioinformatic pipelines originally developed for primary tumors [Rossello et al., 2013, Tso et al., 2014]. Several methods have been developed to facilitate the accurate calling of variants in PDX models. Experimentally, human-specific fluorescence tags can be used to label and isolate human cells prior to DNA extraction [Schneeberger et al., 2016]. Bioinformatically, sequence reads can be aligned to both human and mouse reference genomes, either separately [Conway et al., 2012, Khandelwal et al., 2017] or simultaneously [Bruna et al., 2016], to filter out mouse reads prior to variant calling. Although these approaches greatly improve the reliability of variant calls from PDX models, they entail substantial experimental or bioinformatic burdens. Here we describe a lightweight filtering algorithm that achieves equivalent reliability and can be easily added to standard bioinformatic pipelines.

Many human genes have highly similar paralogous sequences in the genome. Spurious variant calls arising from such paralogs have been recognized as an important source of false positives in the study of rare disease-associated germline variants [Jia et al., 2012, Mandelker et al., 2016, Ng et al., 2010, Zhou et al., 2015]. Similarly, paralogs have led to false positives in the study of cancer, including TUBB in non-small cell lung cancer [Kelley et al., 2001], PIK3CA in hepatocellular carcinoma [Müller et al., 2007, Tanaka et al., 2006], and MLL3 in myelodysplastic syndrome [Bowler et al., 2014]. To address the paralog problem, some variant callers, such as MuTect2 (currently in beta but included in the Genome Analysis Toolkit (GATK; McKenna et al. [2010])), filter clustered variants, which often result from misalignment of paralogous sequences. Many labs also keep lists of suspect genes that tend to suffer from paralog problems and simply ignore any variants called in these genes. These approaches introduce their own biases. Our approach automatically identifies potential spurious calls from paralogs and enables flexible evidence-based filtering.

Here we fully describe and characterize MAPEX (the Mouse And Paralog EXterminator), a BLASTN-based algorithm for filtering variants that was previously introduced by Knudsen et al. [2017]. We also present mapexr, a fast and lightweight implementation in R. We show that, when applied to PDX samples, MAPEX generates calls that are highly similar to other methods, but with less bioinformatic and computational overhead. We also show that, when applied to primary samples, MAPEX effectively filters paralogs while avoiding biases of existing heuristics. MAPEX is thus a useful addition to different tumor variant calling pipelines.

## 2 Approach

### 2.1 Workflow

The MAPEX algorithm is a post-variant-calling filter designed to fit into a standard tumor variant calling pipeline and flag variants which may arise from mis-alignment of mouse reads or from paralogous sequences. The input for MAPEX is a BAM file containing tumor reads aligned to the human reference genome and a variant callset generated from that alignment. MAPEX scores variants by the fraction of variant-supporting reads that align best to the site of the variant when BLASTed against a combined human/mouse reference genome (Figure 1).

**Figure 1:**
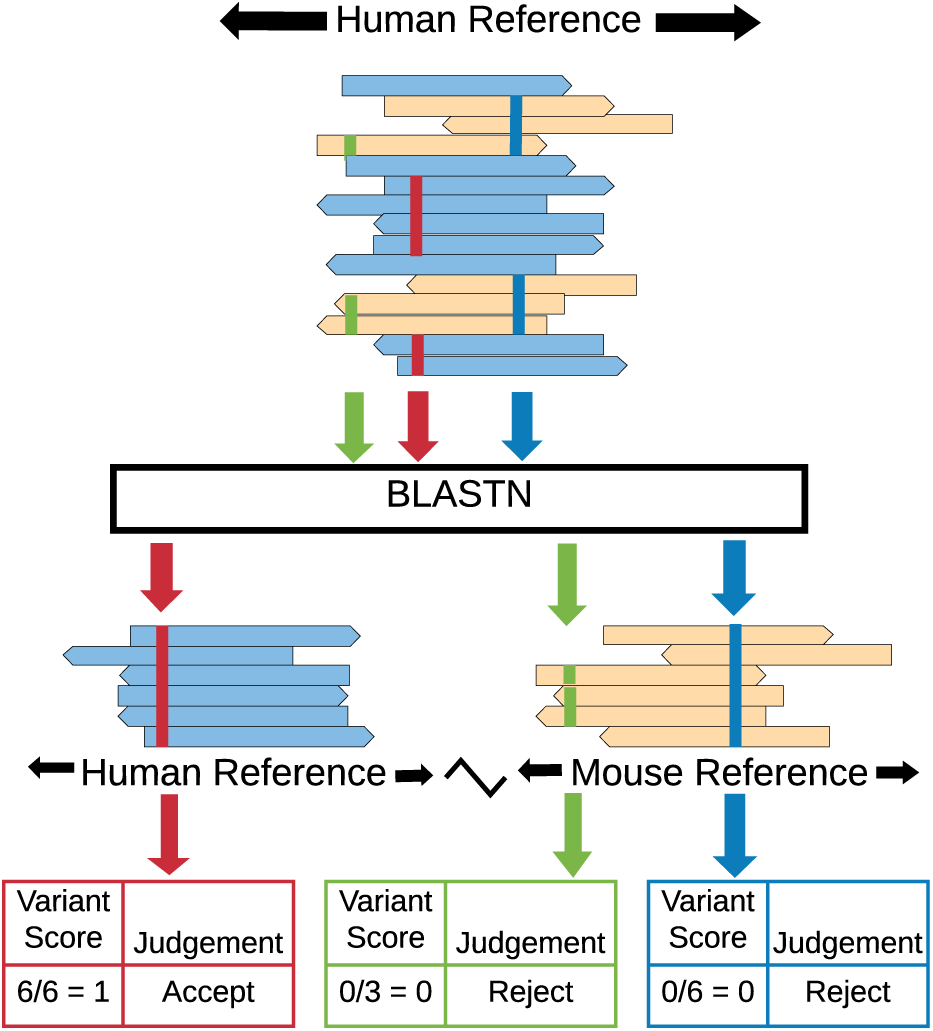
Illustration of MAPEX applied to a PDX sample. MAPEX begins with variants called from tumor reads aligned to the human genome. For each variant (red, blue, and green lines), the supporting reads are BLASTed against the combined human and mouse reference genomes. Variants are then scored by the fraction of supporting reads that align to the called site of the variant in the human genome.

### 2.2 Algorithm

Each read supporting a variant is BLASTed against the appropriate reference genome for the application. For PDX applications, this is the combined human/mouse reference, and for primary tumor applications, this is just the human reference. The best hit for each read is determined by bit score. Reads for which the best hit overlaps the called variant location are classified as “on target” and assigned a score of 1. Reads for which the best hit is a different region of the human genome or a region of the mouse genome are classified as “off target” or “mouse”, respectively, and assigned a score of 0. Reads from genes with close paralogs in the human genome may generate multiple best hits (ties). In this case, the read score is averaged over all best hits, and the read is classified based on the most common result from the best hits. Each variant is then assigned a score that is the average score of all reads supporting that variant and is classified based on the most common classification of the supporting reads.

### 2.3 Implementation

We have implemented the MAPEX algorithm as an R package (mapexr). The package leverages the Bioconductor packages Rsamtools, GenomicAlignments, and GenomicRanges for fast and memory-efficient BAM file handling and read sequence extraction [Lawrence et al., 2013, Morgan et al., 2017]. The package requires a local BLASTN installation and a BLAST database constructed from either a combined human/mouse reference genome or a human reference genome, depending on the application.

## 3 Methods

### 3.1 Samples

To characterize the performance of MAPEX, we used Whole Exome Sequence trimmed fastq reads obtained from pancreatic ductal adenocarcinoma (PDAC) samples described previously by Knudsen et al. [2017] (PDX) and Witkiewicz et al. [2015] (primary). For the PDX analysis, we analyzed a total of 34 PDXs derived from 9 primary tumors, sequenced to mean coverage depth of 124x. For the paralog analysis, we analyzed 93 primary tumors sequenced to a mean coverage depth of 40x.

### 3.2 Alignments and variant callers

All alignments were done using bwa-mem with default parameter settings [Li and Durbin, 2009]. For initial variant calling, we aligned all reads in the samples to the human reference genome GRCh37. We then called variants using MuTect version 1.1.1 [Cibulskis et al., 2013], MuTect2 (as part of the GATK version 3.6, McKenna et al. [2010]), and Varscan 2 [Koboldt et al., 2012], all with default parameter settings. Variants were annotated with Oncotator [Ramos et al., 2015] and the annotation database oncotator v1 ds April052016. For Varscan 2, two PDX samples that yielded millions of variant calls were not processed by MAPEX, to conserve computational time. We considered only non-synonymous single nucleotide variants when comparing between methods. For paralog filtering, we used a conservative variant score cutoff of 0.8.

For comparison with Bruna et al. [2016], we aligned reads to a combined human/mouse reference genome GRCh37/mm9 and called variants using MuTect 1.1.1. We calculated the fraction of mouse contamination using the method described in Bruna et al. [2016]. Briefly, they generated data comparing the fraction of mouse cells in a sample with the fraction of total reads aligned to the mouse portion of a combined reference genome. We used this data to fit a LOESS regression model for contamination fraction vs fraction aligned, and used this to predict mouse contamination based on the fraction of reads aligned to the mouse genome in our samples.

For comparison with bamcmp [Khandelwal et al., 2017], we aligned reads separately to the human and mouse reference genomes and ran bamcmp with default parameters. The output of bamcmp includes alignment files for reads that aligned to only the human reference and that aligned to both references but with a higher human alignment score. We merged these two alignments, performed indel realignment and base score recalibration using the GATK, and used the merged alignment to call variants with Mutect version 1.1.1.

## 4 Results & Discussion

### 4.1 Methodological

MAPEX is a lightweight filtering algorithm that adds little overhead or complexity to existing variant-calling pipelines. The runtime for MAPEX is linear in the number of variants to be filtered. On a 4-core machine, our implementation mapexr, processes roughly 250 variants per minute (Figure S1).

MAPEX has only one tunable parameter, the minimum mapping quality score required for a variant read. The default minimum score is 1, which includes all reads with an unambiguous best mapping. In pipelines in which a minimum mapping quality score is used for variant calling, that score should also be supplied to mapexr, to prevent evaluating reads that were not used by the variant caller. The output from mapexr is an R data frame with four columns – chromosome, start location, variant score, and variant classification – and one row for each variant evaluated. Users may also optionally provide a file path to mapexr which will generate a tab-delimited file with blast results and scores at the read level. The user can choose the variant score threshold used to classify variants as human or mouse derived. Here we use a threshold of 0.5, so that a variant is flagged as spurious if less than half of the supporting reads BLAST as “on target”. In practice, the distribution of variant scores is bimodal and highly concentrated at 0 and 1, so results are insensitive to the exact threshold (Figure S2).

### 4.2 Filtering mouse calls from PDX samples

One important use case for MAPEX is as a post-variant-calling filter for PDX samples that have been aligned to a human reference genome. To test the precision of MAPEX, we compared variant calls from aligning reads to the human reference and filtering with MAPEX to calls from two other methods. The first alternate method is to align reads to a combined human and mouse reference and then call vari-ants [Bruna et al., 2016], which we refer to as the “combined reference” method. The second method is to align reads separately to human and mouse references and call variants using only those reads that align better to the human reference, which is the method implemented in bamcmp [Khandelwal et al., 2017]. For three representative PDX tumors, all three methods yield similar callsets (Figure 2A). The differences are primarily confined to low-frequency variants, and almost all high-frequency variants are called by all three methods (Figure 2B). Across 34 PDX tumors, all three methods yield a similar dramatic reduction in called variants (Figure 2 C).

**Figure 2:**
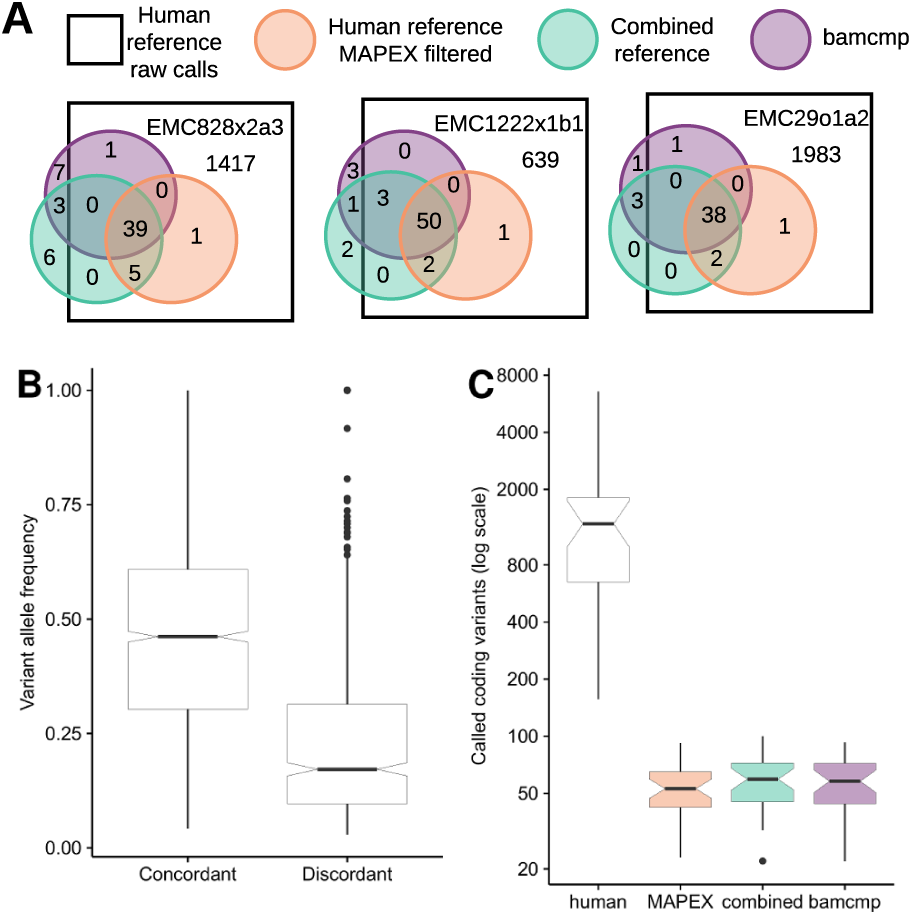
Comparison of MuTect 1.1.1 variants calls between MAPEX, combined reference, and bamcmp methods. A: Detailed breakdown of variant call overlap between the unfiltered human alignment (white square), MAPEX filtered human alignment (red circle), bamcmp filtered human alignment (purple circle) and unfiltered combined alignment (green circle) for representative PDXs created from three different primary tumors. B: Variant allele frequencies for calls that are concordant (n=1663 variants) and discordant (n=552 variants) between the methods. C: Comparison of total calls between the methods, n=34 PDX samples. Boxplots depict 25th and 75th percentile with 1.5×IQR whiskers. Notches are Median ± 1.58×IQR/sqrt(n), and represent rough estimates of 95% confidence interval around the median.

To further validate MAPEX, we compared PDX variant calls before and after filtering to the primary tumor from which the PDX was derived, where mouse contamination is not an issue. Across 34 PDX tumors derived from 9 primaries, MAPEX dramatically enriches PDX calls for variants that were also found in the primary tumor and removes few PDX calls that were found in the primary tumor. Among variants in the PDXs, only 0.3% to 10% called before MAPEX filtering were also found in the primary tumor, but 23% to 90% of variants called after MAPEX filtering were found in the primary tumor (Table S1). This suggests that MAPEX enriches strongly for true variants. Among variants found both in the primary and the PDX before MAPEX filtering, 92% to 100% were retained after filtering (Table S1). This suggests that MAPEX removes few true variants.

To validate the usefulness of MAPEX in practice, we focused on calls within known cancer-associated genes, using the COSMIC database. Among the pancreatic ductal adenocarcinoma (PDAC) samples in COSMIC, 34 genes are mutated in more than 3% of samples. Before filtering with MAPEX, 910 variants were found in these genes among the 34 PDXs we studied. After filtering with MAPEX, only 70 variants were retained. Together, these results suggest that MAPEX removes many false positives, dramatically simplifying variant interpretation. Of particular interest are KRAS, TP53, and SMAD4, which are the most commonly mutated genes in PDAC (Table 1). All of the KRAS mutations filtered by MAPEX are I187V mutants, which result from aligning wild-type mouse KRAS reads to human KRAS, and all 34 PDXs retained the KRAS mutation found in their primary tumor. All of the SMAD4 mutations that were retained by MAPEX in the PDXs also appeared in the corresponding primary tumors. Also of interest is ARID1A, for which the single variant retained by MAPEX was confirmed to appear in the corresponding primary tumor, and none of the filtered variants were present in a corresponding primary tumor.

**Table 1:** Variants detected in PDX samples for important PDAC genes.

| Gene | before MAPEX |  | after MAPEX | COSMIC prevalence |
| --- | --- | --- | --- | --- |
|  | Total variants | Samples with a variant | Total variants |  |
| KRAS | 56 | 34 | 34 | 0.64 |
| TP53 | 9 | 9 | 7 | 0.39 |
| SMAD4 | 5 | 5 | 5 | 0.14 |
| SYNE1 | 3 | 3 | 0 | 0.05 |
| CSMD3 | 96 | 25 | 0 | 0.05 |
| GNAS | 6 | 6 | 6 | 0.05 |
| HMCN1 | 10 | 5 | 0 | 0.04 |
| APC | 12 | 11 | 0 | 0.04 |
| NEB | 31 | 17 | 0 | 0.04 |
| WDFY4 | 6 | 4 | 1 | 0.04 |
| LRP1B | 32 | 18 | 1 | 0.04 |
| ARID1A | 131 | 33 | 1 | 0.04 |

### 4.3 Effects of variant call filters on PDXs

We carried out our primary analyses with the variant caller MuTect 1.1.1, but to test the performance of MAPEX with other variants callers, we also considered MuTect2 and Varscan 2.

If mouse contamination were perfectly filtered, the number of called variants should not depend on the level of mouse contamination. For all three variant callers the number of raw calls was strongly correlated with estimated mouse contamination (Fig. 3A,B,C), although MuTect2 did produce substantially fewer calls overall. After filtering with MAPEX, the numbers of variants called with MuTect 1.1.1 and Mu-Tect2 were not significantly correlated with the level of mouse contamination (Fig3D&E). On the other hand, the number of variants called with Varscan 2 was correlated with mouse contamination (Fig. 3F), suggesting that MAPEX is not eliminating all spurious calls.

Importantly, as a post-variant-calling filter, MAPEX can not evaluate variants that were not initially called. Filters implemented with a variant caller, generally designed to improve results from primary tumors, can cause problems when using MAPEX. For example, MuTect2 applies a clustered event filter designed to reduce the number of false-positive variant calls due to mis-alignment of highly paralogous sequences. In regions of high similarity between mouse and human, this filter can remove true variants. For instance, Figure 4 shows the result of aligning a PDX with modest mouse contamination to the human reference for a small portion of the KRAS oncogene. MuTect 1.1.1 and Varscan 2 both called three variants at this locus, and MAPEX correctly rejected the two spurious variants arising from mouse contamination and retained the true G12D variant. MuTect2 fails to call any of these variants, because they are filtered as likely homologous mapping events, so MAPEX does not see and cannot retainthe true G12D variant. In our PDX samples, we found instances of the clustered event filter removing true variants from other PDAC oncogenes, including SMAD4 and TP53.

**Figure 3:**
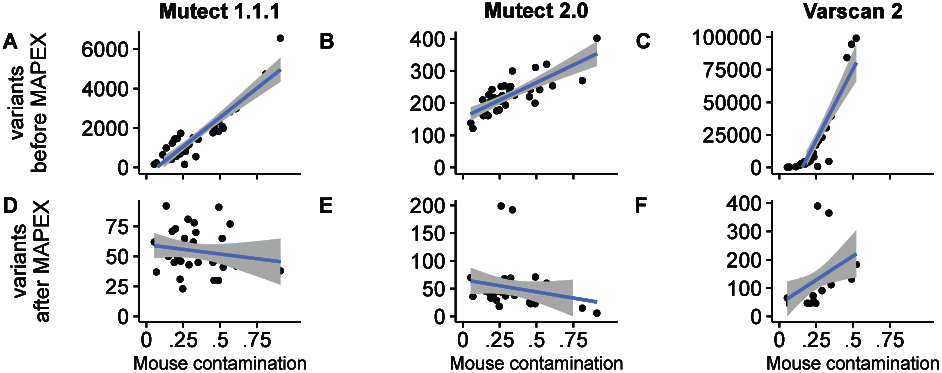
Effects of variant caller on analyzing xenograft samples with MAPEX. A,B,C: For all three calling algorithms and 34 xenograft samples (black dots) the number of raw variants called was strongly dependent on estimated mouse contamination. D,E,F: After filtering with MAPEX, the number of calls was independent of mouse contamination for MuTect 1.1.1 and MuTect2, but not for Varscan 2. Blue lines show linear regressions and shading denotes 95% confidence intervals.

**Figure 4:**
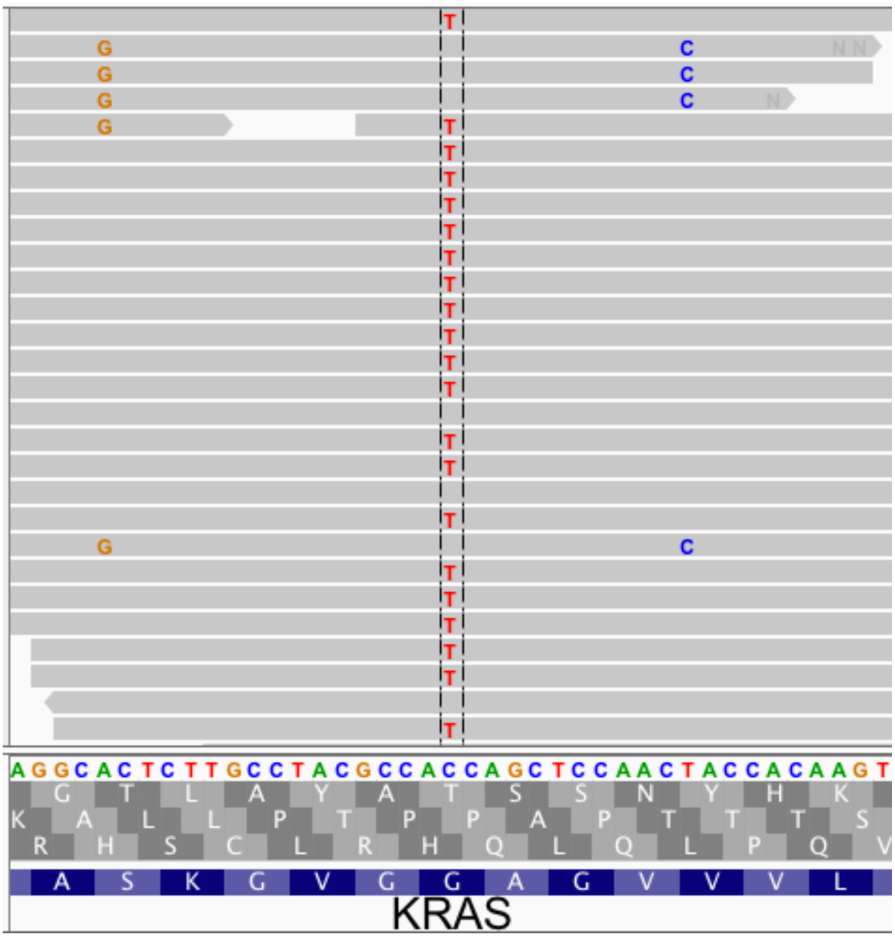
This Integrative Genomics Viewer [Thorvaldsdottir et al., 2013] window covers a portion of the human KRAS gene. The C>T variant is the classic KRAS G12D mutation that appears in many PDAC tumors. The A>G and T>C variants both result from aligning wild-type mouse reads to the human sequence. When used with MuTect 1.1.1 or Varscan 2, MAPEX correctly retains only the G12D variant. MuTect2, however, filters all three variants, so the G12D variant cannot be retained.

Overall, the performance of MAPEX does not depend sensitively on the variant caller used, but callers can introduce specific biases. In particular, the default parameters for Varscan 2 yield high sensitivity but low specificity. When Varscan 2 is applied to PDX samples with mouse contamination, MAPEX thus does not filter out all spurious calls. As such, we recommend that users of Varscan 2 be cautious when calling PDX samples and perhaps apply additional post-calling filters. By contrast, the default parameters for MuTect2 yield much higher specificity, but at the cost of sensitivity in the PDX context. Currently, the clustered event filter cannot be disabled in MuTect2. We thus advise that users pairing MAPEX with MuTect2 be cautious when interpreting callsets from PDX samples in genes with high similarity between human and mouse.

### 4.4 Flagging potential false positives resulting from paralogous sequences

In addition to removing mouse contamination from PDX samples, MAPEX can also filter potential paralogs in primary samples. Across 93 PDAC primary tumors, a mean of 11% of total variant calls were flagged by MAPEX as potential paralogs, with a range of 2-33%. The genes in which variants were most frequently flagged as potentially arising from paralogous sequences include members of large gene families, such as mucins, zinc-finger nucleases, and the PRAME family (Table 2). Variants in citrate synthase (CS) were also frequently flagged (Table 2). Citrate synthase has a known pseudogene NCBI: LOC440514, which was responsible for all of the spurious calls. We called variants with MuTect 1.1.1 and filtered with MAPEX, but MuTect2 includes new clustered event and read-mapping quality filters to prevent calling variants caused by paralogs. Using MAPEX yielded call sets that were identical with MuTect2 for all the genes in Table 2, with the exception of MUC12 and MUC5B, which differed by 3 variants. MAPEX can thus be efficiently and confidently used to remove variants that likely arise from paralogous sequences.

**Table 2:** Top genes for which MAPEX flagged variants as potentially arising from paralogs.

| Gene | Variants<br>flagged | Samples with<br>a flagged variant |
| --- | --- | --- |
| ZNF814 | 15 | 15 |
| CS | 12 | 7 |
| IGFN1 | 8 | 6 |
| KMT2C | 7 | 7 |
| FRG1 | 6 | 6 |
| LILRB3 | 6 | 6 |
| MUC12 | 6 | 6 |
| RGPD3 | 6 | 6 |
| USP6 | 6 | 3 |
| FCGBP | 5 | 4 |
| MUC5B | 5 | 5 |
| NBPF1 | 5 | 3 |
| PRAMEF11 | 5 | 4 |
| PRB4 | 5 | 3 |
| RGPD8 | 5 | 4 |

## 5 Conclusion

Genome sequencing is an increasingly important tool in cancer research, but spurious variant calls remain a challenge. MAPEX is an algorithm designed to filter spurious variants caused by mouse reads in patient-derived xenografts (PDXs) and caused by paralogous sequences in primary tumors. We showed that MAPEX is as sensitive and specific as more computationally intensive methods for calling variants from PDX tumors. We also showed that MAPEX successfully flags variant calls in potentially problematic gene families in primary tumors. Our implementation, mapexr, fits cleanly into standard tumor variant-calling pipelines and runs quickly on modern desktop computers. MAPEX is thus a potentially useful new component for many tumor variant-calling pipelines.

---

**Figure S1:**
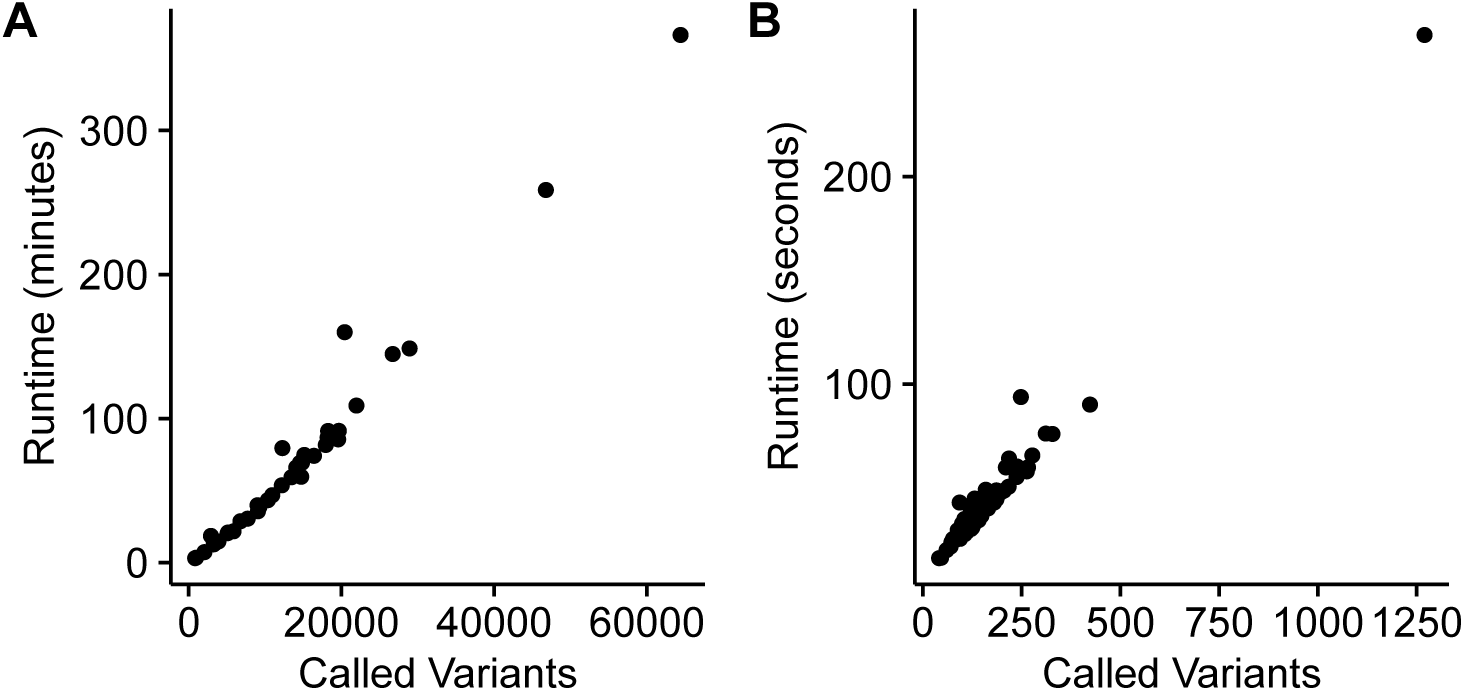
mapexr timing for A: xenografts and B: primary tumors. Shown are results from running on 4 cores and filtering all MuTect 1.1.1 calls for each sample. Run time is linear in the number of input variants, roughly one minute per 250 variants. One strategy for reducing run time is to first filter to keep only variants of interest, such as non-synonymous coding variants.

**Figure S2:**
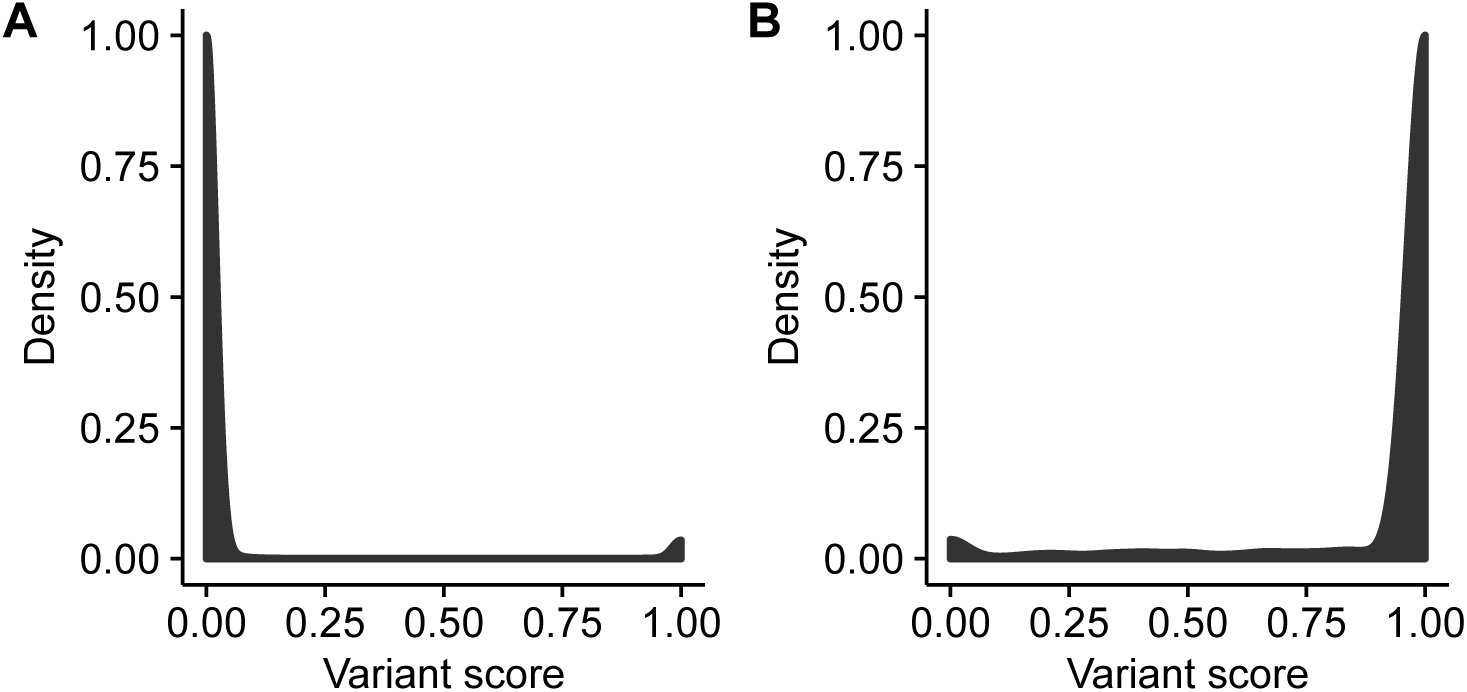
Distribution of variant scores from MuTect 1.1.1 over all A: PDX samples and B: Primary samples.

**Table S1:**
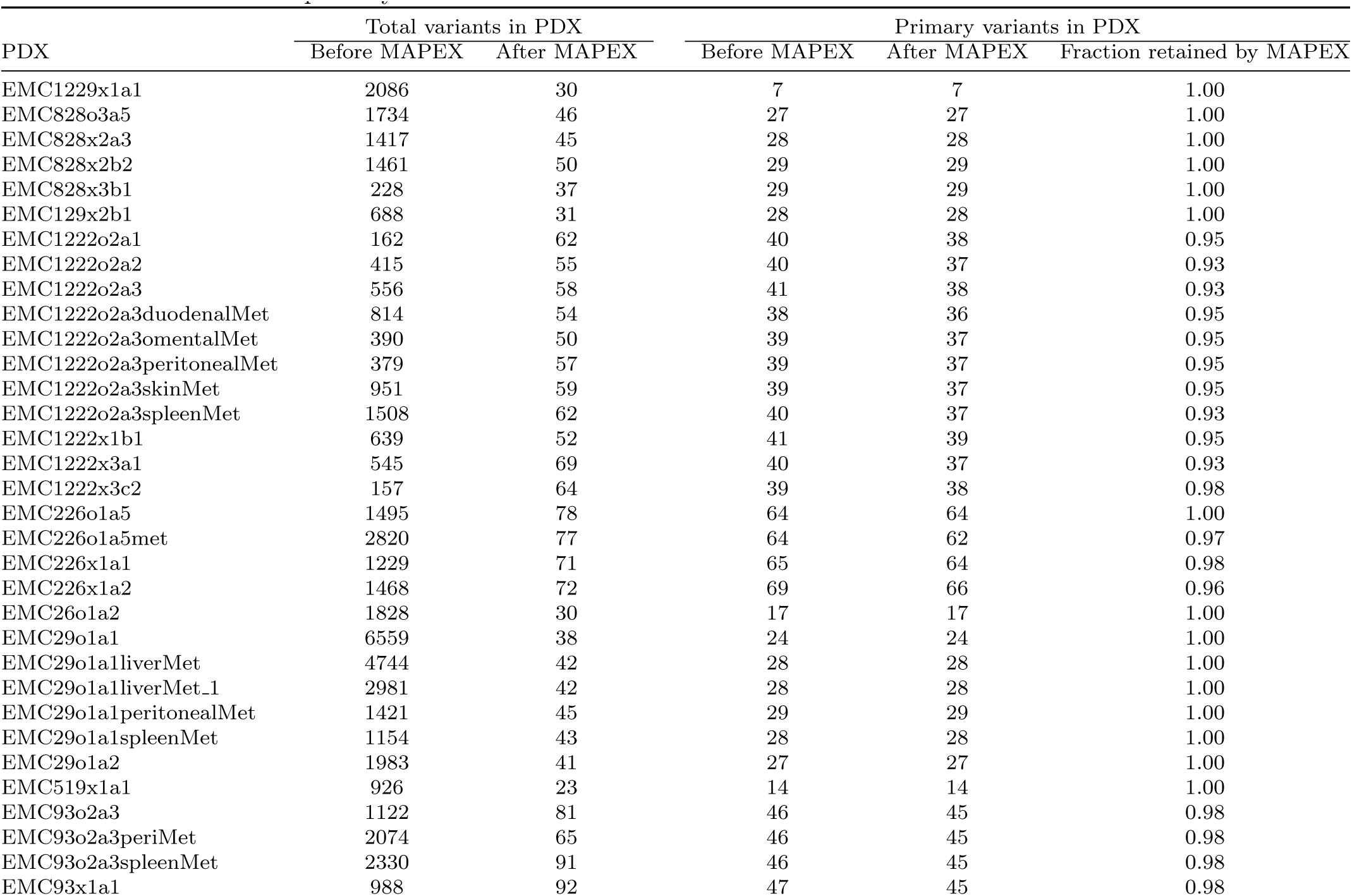
MAPEX removes many potentially spurious variants and retains almost all likely real variants that are also found in the primary.

